# Diversity *Between* but Not *Within* Microbial Communities Increases With the Diversity of Supplied Nutrients

**DOI:** 10.1101/2025.05.10.653264

**Authors:** Arthur Newbury, Angus Buckling

**Affiliations:** Environment and Sustainability Institute & Department of Ecology & Conservation, College of Life and Environmental Sciences, Biosciences, University of Exeter Penryn Cornwall TR10 9FE, UK

## Abstract

Greater resource diversity within a habitat is typically expected to support greater biodiversity in the resident ecological community (*α*-diversity). It may also make community composition less deterministic since a wider range of species have a chance establishing themselves and affecting subsequent community assembly processes. However, the relationship between microbial diversity and the diversity of resources exogenously supplied is complicated by cross-feeding between microbes, since much of the resource diversity encountered will be generated by the community. Here, we assembled microbial communities in simple growth media (where cross-feeding will be important) with 1 to 3 carbon sources and asked how increasing nutrient diversity affects diversity within and between communities. While there was (i) no clear difference in *α*-diversity, increasing the number of carbon sources led to communities that varied (ii) more within and (iii) less between the different abiotic conditions. (i) is at odds with classical ecological theory, but fits predictions from a consumer-resource model that accounts for cross-feeding. (ii) shows an increase in the stochasticity of community assembly. (iii) demonstrates strong abiotic selection in the single carbon source communities. We then went on to show that this is leads to phylogenetic clustering (co-occurrence of more closely related strains).

## Introduction

How multiple competing species coexist within ecological communities is a central question in ecology (Hardin 1960). One logical assumption is that the availability of a wider variety of ecological niches should support greater biodiversity. This is illustrated mathematically in the consumer resource framework of McArthur (MacArthur 1970), with Tillman showing that species diversity depends on having a diversity of resources and a diversity of resource requirements (Tilman 1982). For microbial communities such consumer-resource relationships are highly non-linear - diversity begets diversity as bacteria produce secondary metabolites, which are fed upon by neighbouring microbes (Duan et al. 2009; Leventhal et al. 2018). This may be one of the reasons microbial communities are so much more diverse than other types of ecological communities. As a consequence, it is difficult to predict the effects of increasing the diversity of nutrients not produced by the microbes themselves.

Recent work has shown that increasing nutrient diversity affects a very small increase in the biodiversity of experimental microbial communities e.g. a non statistically significant increase of approximately one additional microbial strain going from a resource diversity of 1 to 32 (Pacheco, Osborne, and Segrè 2021) or a linear increase of one strain per additional carbon source (Dal Bello et al. 2021). A potential reason for this is that much of the nutrient diversity needed to sustain a microbial community is provided by the microbes within the community (Dal Bello et al. 2021).

While the direction of the effect on *α*-diversity is at least consistent with predictions, it is less obvious how *β*-diversity (differences between communities) will be affected by the combination resource diversity supplied from outside of the communities and secondary metabolites produced by the communities. Having a wider range of nutrients supplied may increase the pool of potential core taxa, leading to less determinism in community assembly. However, this effect could plausibly be either dampened or exacerbated by cross-feeding interactions. As with *α*-diversity, if secondary metabolites account for much of the resource diversity then an increase in the diversity of externally supplied resources may have a limited effect. On the other hand, reduced determinism in the core strains could have large impacts on which secondary consumers establish in the community (Leventhal et al. 2018).

Here we combine mathematical modelling and experimental ecology to investigate the impacts of nutrient diversity on both *α* and *β*-diversity. Consumer-resource models and microbial communities cultured on one or a combination of carbon sources are used to highlight the potentially important role of cross-feeding (microbes consuming metabolites secreted by other community members) in subverting the expected relationship between nutrient diversity and biodiversity. This is followed by a more detailed investigation of community assembly, identifying the role of abiotic selection in shaping the phylogenetic structure of metacommunities.

## Results

### α-diversity

The same starting stock of 46 bacterial isolates were used to seed 84 minimal media microcosms - 12 replicate microcosms for each carbon source (arabinose, glucose and leucine) as well as each combination of multiple (2 or 3) carbon sources. In order to determine the effects of resource diversity on biodiversity and community assembly, we split the microcosms into two groups - single and multiple carbon sources. *α*-diversity was measured as the number of strains surviving in each microcosm, the evenness of strain abundances (Pielou’s index) and the phylogenetic diversity of each community (Faith’s index). Separate Bayesian models were fit for each diversity metric to determine the effect of having multiple carbon sources on *α*-diversity in general as well as the effect relative to each single carbon source (see methods for model selection and final model structures). In all there was little evidence of any effect of resource diversity on *α*-diversity, with nutrient diversity having an estimated effect of -0.015 (CI[-0.414, 0.394], PD = 0.566) on log strain richness, 0.092 (CI[-0.554, 0.782], PD = 0.68) on log-odds strain evenness and 0.037 (CI[-0.04, 0.114], PD = 0.83) n log phylogenetic diversity (Figure 1 a-i). The only exception was that the leucine communities were less even than the multiple carbon source communities by 0.353 (CI[0.172, 0.542], PD > 0.999) (Figure 1 f), although they were also less even than the other single carbon source communities (Figure 1 d).

**Figure 1.**
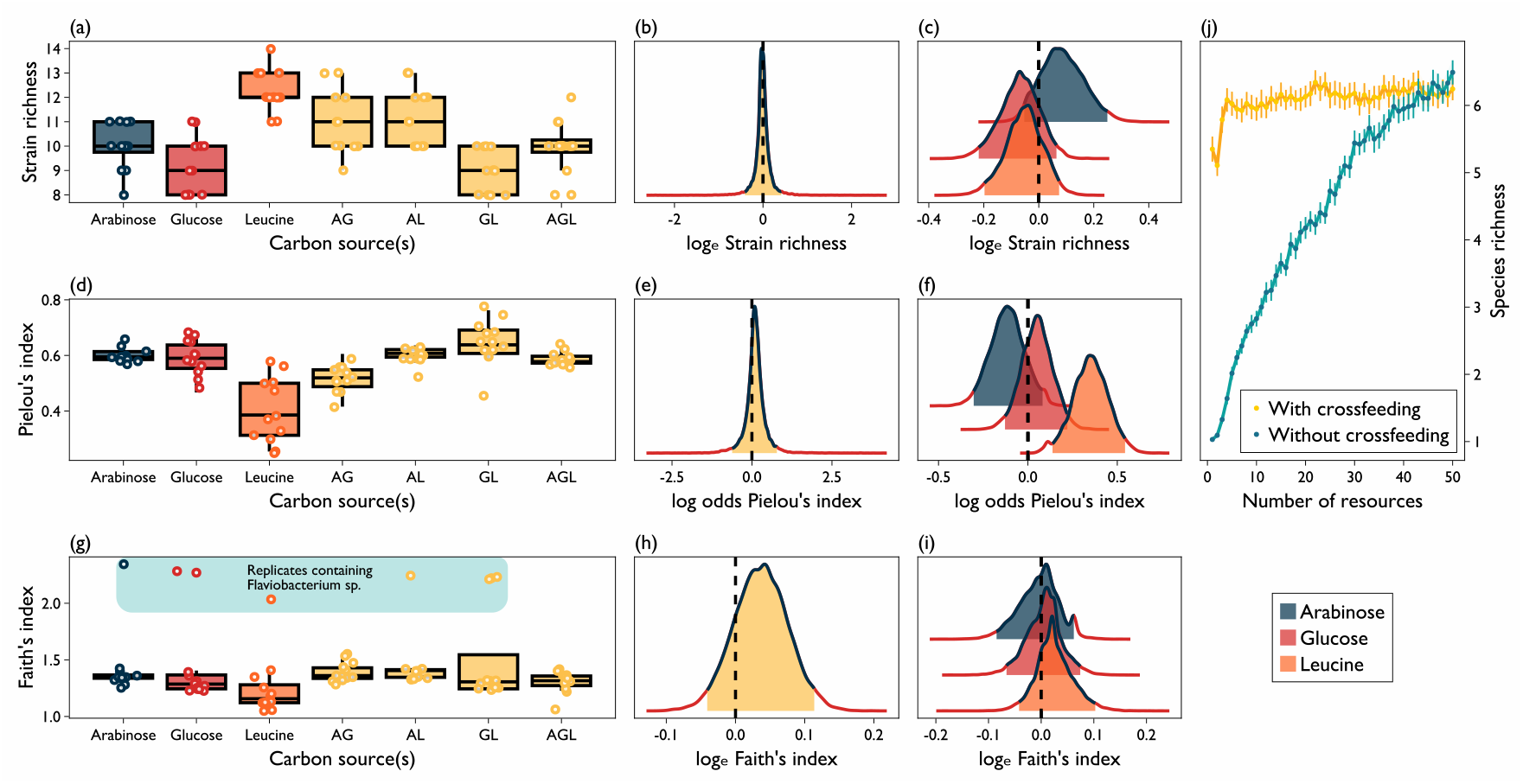
*α*-diversity. (a-i) Raw data (a, d & g) and estimated effect of having multiple carbon sources globally (b, e & h) and per single carbon source (c, f & i) for strain richness, evenness (Pielou’s index) and phylogenetic diversity (Faith’s index) respectively. In panel (g) we highlight the Faith’s index outliers owing to the presence of a single rare isolate *Flavobacterium sp.* which was distantly related to the other strains. Boxes indicate interquartile range, with whiskers denoting the range of the data. (j) Consumer resource model results, showing the effect of resource diversity on species richness with or without the ability for consumers to excrete resources. Error bars show the standard error of the mean.

The above result is seemingly at odds with standard ecological theory, as we would expect that having a variety of resources to specialise on would reduce niche overlap between strains and allow greater coexistence. However, this may not apply to microbial communities where much of the nutrient diversity comes from secondary metabolites released by the microbes themselves. We explored the possible impact of this using a consumer-resource model. The model which follows the same basic structure used in recent publications (Goldford et al. 2018; Marsland III, Cui, and Mehta 2020; Pacheco, Osborne, and Segrè 2021) was used to measure the relationship between the number of resources (continuously supplied) to number of extant consumer species at equilibrium, with no cross-feeding. This was repeated for the case where the consumer species each converted those primary resources into a range of secondary resources which were then *leaked* into the simulated environment to be consumed by the rest of the consumer community. The result of allowing cross-feeding was greatly increased consumer diversity at lower levels of primary resource diversity and thus a far flatter response to increasing resource diversity.

We assessed whether cross-feeding was important in our experimental communities by growing each strain in isolation in minimal media with each carbon source and in the mix of all 3 carbon sources, with and without the addition of spent media supernatant from propagating the entire community together in the given growth media. In each carbon source, only a subset of the 46 isolates showed any detectable growth after 24 hours: 22, 13 and 7 for arabinose, glucose and leucine, respectively. Of these only a subset were able to grow without community spent media: 6 of 22, of 8 of 13 and 5 of 7 for arabinose, glucose and leucine respectively (Figure 2). This confirms that many of the isolates used can benefit from the by-products of others. However, our results also demonstrate that these short-term growth assays were not a good predictor of establishment within experimental communities, since many isolates that persisted in communities did not exhibit appreciable population growth in monoculture within 24 hours (black points in Figure 2 a). Furthermore, there was more detectable growth in the mixture of all 3 carbon sources, 32 strains in all, of which only 12 grew without supernatant. It is unsurprising that more strains were able to grow in the media with more carbon sources, but this fits with the expectation that we would see more diversity (*α, β* and *γ*) with multiple carbon sources.

**Figure 2.**
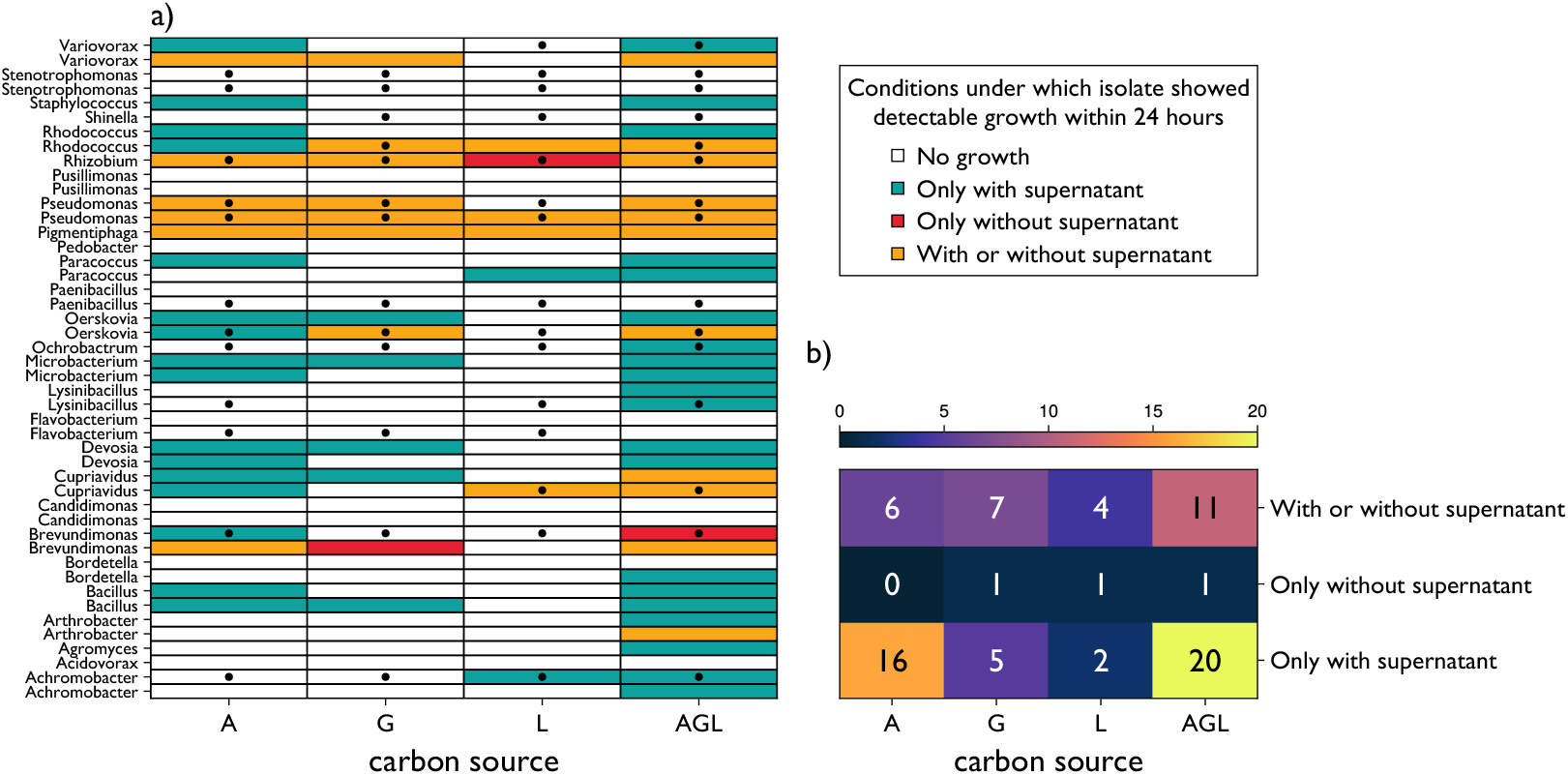
Monoculture growth. a) Conditions under which isolates grew noticeably within 24 hours, with black points signifying that a given isolate was detected at the end of the main experiment at least once in the corresponding carbon source treatment. b) Total number of isolates for which growth was detected under the given conditions. Carbon sources are denoted by initials, with AGL representing the mixture of all 3.

### β-diversity

*β*-diversity (the dissimilarity between communities) may increase with nutrient diversity, since a wider range of taxa can feasibly survive, opening up the possibility for possible community structures to emerge. However, given the *α*-diversity results, it is unclear if cross-feeding (or some other process) will overturn or nullify the expected result. Thus we contrasted the *β*-diversity exhibited by single carbon source communities with that of multiple carbon source communities. The hypothesis that increased resource diversity should lead to increased *α*-diversity rests on the assumption that the available resources do not all select for the same members of the species pool. Therefore, we also measured the dissimilarity between pairs of communities cultured on different carbon sources (or different mixtures in the multiple carbon sources case) to confirm that communities cultured in different media are more dissimilar than are pairs of communities cultured on the same carbon source. While our main focus is on the presence vs absence of strains, dissimilarity in terms of relative abundance is another sign of the abiotic environment having a selective effect. Thus, we measured Bray-Curtis dissimilarity (based on proportion data) and weighted UniFrac, alongside Jaccard’s index and unweighted UniFrac. For both single and multiple carbon source conditions we estimated the dissimilarity between pairs of communities from either the same or different abiotic environments using a mixed model to account for non-independence of data in dissimilarity matrices.

In all cases dissimilarity was significantly greater between than within abiotic conditions (PD > 0.999 in each case), demonstrating that the carbon sources used do indeed select for different strains. Additionally, the difference between abiotic environments was greater in the single carbon source condition (PD > 0.999 in each case), indicating that swapping 1 single carbon source for another has a larger impact on community composition than changing only 1 of 2 carbon sources e.g. going from arabinose to glucose vs going from arabinose + leucine to glucose + leucine. However, in terms of within carbon source *β*-diversity, whether or not this was higher with multiple carbon sources depended on how *β*-diversity was measured. For Jaccard’s index there was an increase in *β*-diversity with multiple carbon sources of 0.039 (CI[0.019, 0.06], PD > 0.999). Whereas, for unweighted UniFrac there was a decrease in *β*-diversity with multiple carbon sources of 0.038 (CI[0.021, 0.056], PD > 0.999). Thus, while resource diversity did lead to reduced determinism in terms of which strains survived in a community, in our data set this did not hold when accounting for the relatedness between strains. However, this discrepancy was driven by a single rare isolate *Flavobacterium sp.* which had large leverage in our UniFrac analyses due to being distantly related to the other strains. Results excluding this strain were in line with Jaccard’s index results (see supplementary Figure 1).

Given the *α*-diversity results, it is obvious that multiple carbon source communities did not contain the combined taxa of the respective single carbon source communities. A clear example of the non-additive relationship between nutrient diversity and strain richness (Figure 1 a) is *Cuprivadious sp.* which was not present in arabinose or glucose communities, or in the mix of all three carbon sources, but was found in 5 out of 12 of the arabinose-glucose mix communities (Figure 3 a). Another example is *Ochrobactrum sp.*, which was present in glucose and leucine communities, but not in the mixture of both.

**Figure 3.**
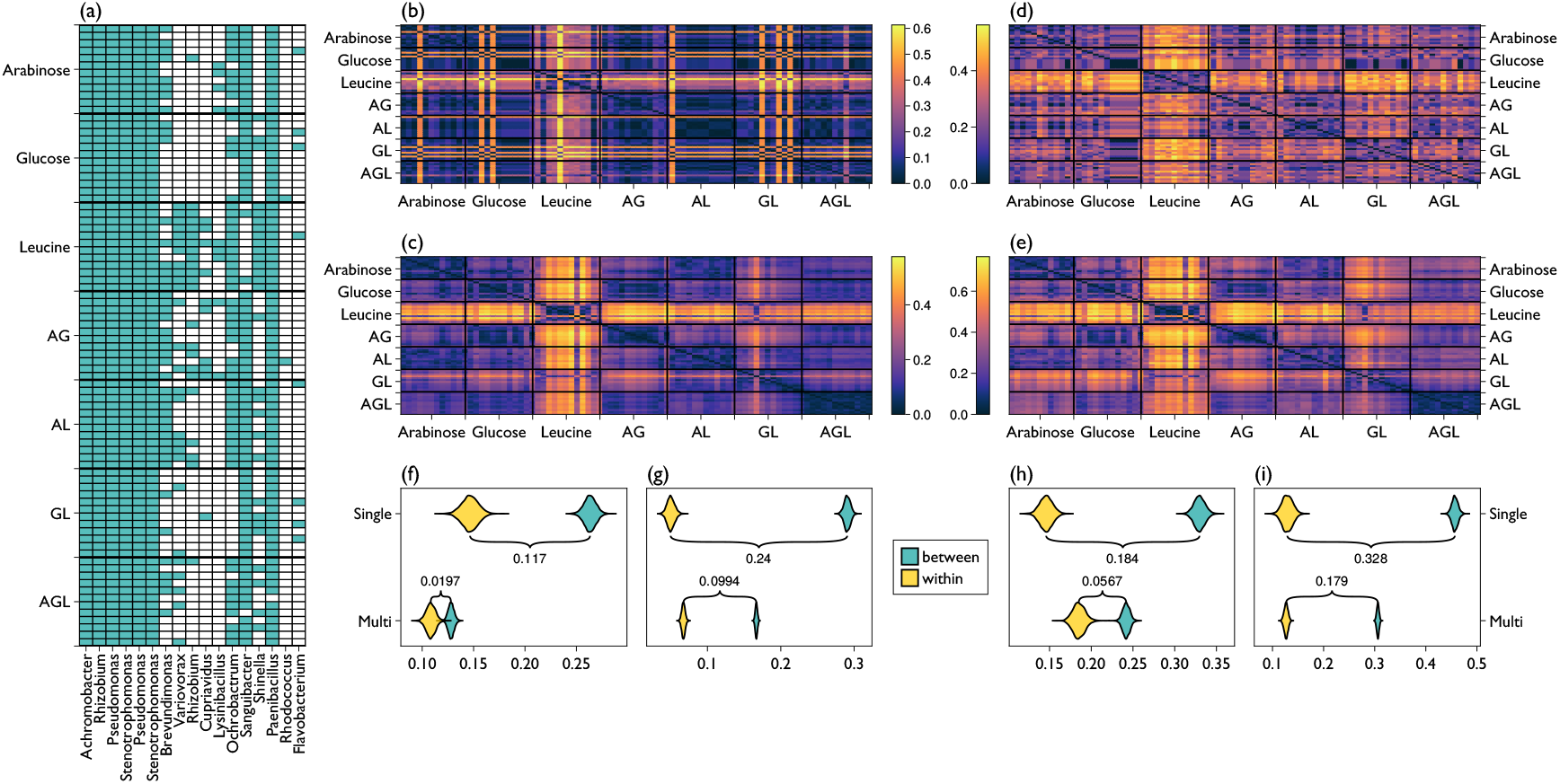
(a) Presence/absence of strains in each microcosm. (b-e) Unweighted and weighted UniFrac, Jaccard’s and Bray-Curtis dissimilarity respectively. (f-i) Posterior estimates of dissimilarity both between and within abiotic treatments of both single and multiple carbon source conditions - Unweighted and weighted UniFrac, Jaccard’s and Bray-Curtis dissimilarity respectively.

### co-occurrence and relatedness

As a consequence of habitat filtering, single carbon source minimal media selects for closely related taxa (phylogenetic clustering) when comparing between different treatments (Goldford et al. 2018). However, it is unclear if these more closely related strains tend to co-occur within individual communities. Classical ecological theory would predict closely related strains to exclude each other within these simple experimental microcosms (Hardin 1960). To determine whether competition between closely related strains is sufficient to disrupt the treatment level phylogenetic clustering at the level of individual communities we analysed the relationship between co-occurrence affinity (Mainali et al. 2022) and phylogenetic distance for pairs of strains separately for single and multiple carbon source conditions using a model developed in (Newbury 2024). Positive co-occurrence affinity values indicate that a pair of strains tend to co-occur more frequently than expected by chance, given the prevalence of each strain. The relationship between co-occurrence and phylogenetic distance was measured separately for single and multiple carbon source groups. Co-occurrence affinity decreased with phylogenetic distance for single carbon source communities, with a regression coefficient () of -2.835 (CI[-6.251, 0.556], PD = 0.946), whereas in the multiple carbon source communities there was no obvious trend: = -0.398 (CI[-3.355, 2.557], PD = 0.603) (Figure 4).

**Figure 4.**
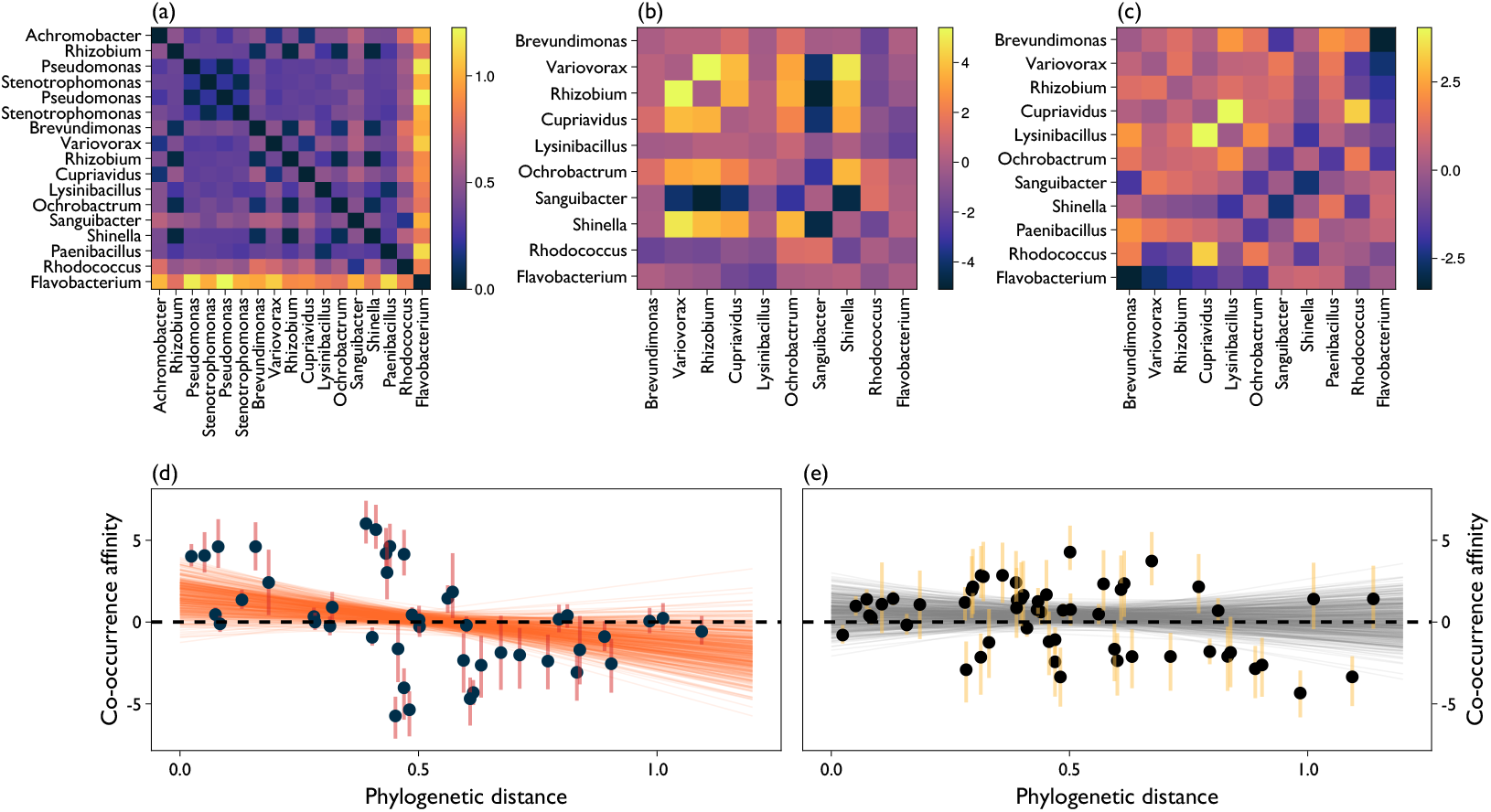
Co-occurrence and relatedness of strains. (a) Phylogenetic distance between all strains detected at the end of the experiment. (b-c) Maximum a posteriori (MAP) estimates of co-occurrence affinity between all analysable pairs of strains (i.e., where neither strain is either always present or always absent) in the single (b) and multiple (c) carbon source conditions. (d-e) Relationship between phylogenetic distance and co-occurrence under single (b) and multiple (c) carbon source conditions. Error bars denote interquartile range of posterior estimates of co-occurrence affinity. Line plots are draws from posterior estimates of linear relationship between phylogenetic distance and co-occurrence affinity.

It is possible however that competition within genera led to the exclusion of particular strains from all microcosms (Figure 2 a). This would imply a hump-shaped relationship between phylogenetic distance and co-occurrence. However, this complete exclusion of very closely related taxa would not be seen by our model as it can only be applied where there is variation within the data set i.e., strains that are present in some but not all communities. *Pseudomonas* and *Stenotrophomonas* each had 2 strains present in every single community (Figure 3 a) and these were the only cases of of more than 1 strain being present from a single genus. In order to determine whether this number was smaller than we would expect by chance (implying competitive exclusion within genera) we randomly sampled (without replacement) from the 46 isolates with a number of draws equal to the strain richness of each community 100,000 times for a total of 8,400,000 randomly assembled communities. We found that the probability of getting 2 genera with multiple strains was 0.185 and the likelihood of having more than 2 was 0.039. Thus, we actually see more co-occurrence within genera than expected by chance. We also repeated this procedure but focused on the *γ*-diversity of the whole experiment - 8,400,000 randomly assembled communities with 17 draws each time. Here, 2 was the most likely result, with a probability of 0.326 and the likelihood of having more than 2 genera with multiple strains was 0.489, approximately even odds. Again, no evidence that competition within genera had an overly large impact on community or metacommunity composition.

## Discussion

We have shown that for a community of bacteria, increasing nutrient diversity will not necessarily increase biodiversity even if it affects community composition. A consumer-resource model accounting for the widespread cross-feeding in microbial communities predicts that this one change could account for the disconnect between the diversity of nutrients supplied and the biodiversity of the community. However, nutrient diversity did predictably increase *β*-diversity. Furthermore, the negative relationship we observed between phylogenetic distance and co-occurrence affinity was minimised in the multiple carbon sources treatment, where the abiotic differences between microcosms was reduced. Thus, both the identities and number of carbon sources did impact community assembly, just not such that it significantly impacted *α*-diversity.

Microbial diversity is typically far greater than the diversity of other organisms within a given area (Labouyrie et al. 2023; Consortium 2012). Here, given the 6 week length of this experiment, that between 8 and 14 strains in a randomly assembled community could coexist in an extremely alien and very simple habitat is noteworthy (similar studies have utilised shorter time scales of 1 week (Dal Bello et al. 2021) or low numbers of strains (Pacheco, Osborne, and Segrè 2021)). A parsimonious explanation borne out in the modelling results presented here is that the widespread production of secondary metabolites means that microbial communities do not require a great diversity of primary nutrients in order maintain coexistence. That is - the key to microbial diversity may be the levelling of the food chain/web - that each microbe provides niche space for another. The diversity enhancing properties of cross-feeding may be particularly pronounced in spatially structured communities, where the local microbial populations determine local selective conditions through the secretion of secondary metabolites (Newbury, Kuijper, and Buckling 2023). However, here (as in previous studies (Wawrik et al. 2005; Goldford et al. 2018)) carbon source did predict community composition (Figure 3). Thus, primary resources are not irrelevant, but rather determine the context-dependent hierarchical structure of microbial trophic relationships. The effects of habitat filtering were in fact far more prominent in the structuring of communities than was competition between closely related strains.

In our experimental microcosms, closely related strains tended to co-occur with greater frequency than more distantly related ones. Insights primarily from plant community ecology illustrate that the co-occurrence of related taxa (phylogenetic clustering, as opposed to phylogenetic over-dispersion) results from the interplay between the relative importance of habitat-filtering vs competitive exclusion and the degree of trait conservation in evolution (Webb et al. 2002). Since microbial communities converge on coarse grained patterns based on the available nutrients, and since the secretion of secondary metabolites allows for high diversity with even only a single supplied carbon source (Goldford et al. 2018), microbial communities may be more likely to exhibit phylogenetic clustering than communities of animals or plants. This is because the predictability of assemblages based on nutrient source suggests strong habitat filtering (leading to closely related taxa favouring the same locations), while the stabilising effect of cross-feeding may allow these closely related taxa to coexist rather than exclude each other. Indeed, a positive relationship between relatedness and co-occurrence has been observed in natural microbial communities (Kamneva 2017; Goberna et al. 2019). However, understanding the mechanisms that lead to the observed patterns in community assembly requires assembling communities under controlled conditions, since as well as habitat filtering, phylogenetic clustering can occur due to evolutionary diversification within a given clade, with limited dispersal to distant geographic regions (Webb et al. 2002). Here, by seeding replicate communities with the same starting stock of bacteria we are able to rule this process out. The causal effect of habitat filtering is further illustrated by the fact that the positive relationship between relatedness and co-occurrence is only apparent amongst the single carbon source communities (Figure 4) where the treatments create larger differences in community composition (Figure 3 h).

Taken together, our results highlight the importance of considering microbial communities as self-reinforcing systems with relatively less reliance on externally produced niches to support biodiversity. Though the specific make-up of the community will be strongly influenced by abiotic factors, the diversity enhancing properties of microbial communities allow even closely related strains to coexist in simple habitats with limited diversity in supplied nutrients. This latter is an important consideration for microbiome engineering, where it may be desirable to culture a set of closely related but functionally distinct strains.

## Methods

### Consumer-resource model

To investigate how resource diversity and cross-feeding interact to affect biodiversity, we employed a consumer resource model implemented in MiCRM.jl (Clegg 2022) which we solved numerically using Tsitouras’ 5/4 Runge-Kutta method (Tsitouras 2011) implemented in DifferentialEquations.jl (Rackauckas and Nie 2017). The model takes the form

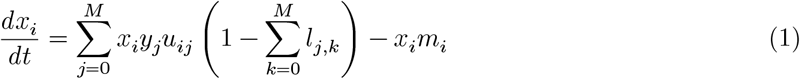

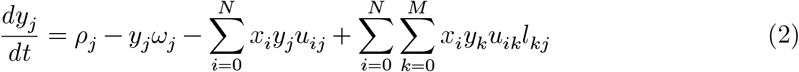

where *x*_*i*_is the biomass of the *i*th consumer, *y*_*j*_is the mass of the *j*th resource, *N* in the number of consumer populations, *M* is the number of resource types, *u*_*ij*_is the rate of uptake of the *j*th resource by the *i*th consumer, *ρ*_*j*_and *ω*_*j*_are the inflow and outflow rates of the *j*th resource, *m*_*i*_is the loss rate of the *i*th consumer and *l*_*jk*_is the proportion of the uptake of the *j*th resource that is leaked to the *k*th resource. To measure the effect of resource diversity on consumer diversity without cross-feeding, we ran numerical simulations with 30 consumers, between 1 and 50 resources with inflow rates *ρ* equal to 1/*N*, outflow rates *ω* drawn from uniform distribution between 0 and 0.001/*N*, all *m*_*i*_= 1 and no leakage of resources. To measure the same effect in the presence of cross-feeding, we ran additional simulations as above but with an additional 50 resources that can only be produced via leakage from the *primary* resources with each corresponding leakage rate *l* drawn from uniform distribution between 0 and 0.03. After running for 2000 time steps (found to be long enough for the system to reach equilibrium) diversity was calculated as the number of consumer populations with biomass greater than an extinction threshold of 0.001.

### Microbial communities

46 bacterial isolates (previously isolated by Hesse et al. (2018)) from 27 genera were inoculated at ≈ equal densities into 1X M9 buffer to form a multispecies stock solution. From this 60 µl was used to inoculate each replicate microcosm. Communities were cultured at 28°C in 25 mL glass microcosms containing 6 mL M9 minimal media with 0.07 C-mol/L of either arabinose, glucose or leucine, one of the 3 possible pairwise combinations (each carbon source contributing 0.035 C-mol/L) or a mix of all 3 (each carbon source contributing 0.023 C-mol/L) - n = 12 for each nutrient treatment. 60 µl of culture was transferred to a fresh microcosm every 7 days for a total of 6 weeks growth.

### DNA extraction and sequencing

Following biofilm disruption via the vortex—sonicate—vortex method (Mandakhalikar et al. 2018), genomic DNA was extracted using the Qiagen blood and tissue kit (Qiagen, Hilden, Germany) following the manufacturers standard protocol. Library preparation via 2-step PCR amplification of V4 region of 16S rRNA, and sequencing performed on an Illumina MiSeq v2 (Paired-end, 2×250 bp) were carried out at the Liverpool centre for Genomic Research.

### Bioinformatics

Raw reads were trimmed for the presence of Illumina adapter sequences, using Cutadapt version 1.2.1 (Martin 2011) to trim the 3’ end of all reads matching the adapter sequence for at least 3 bp. Further trimming was implemented using Sickle version 1.200 (https://github.com/najoshi/) with a minimum window quality score of 20. Reads were further trimmed, filtered, denoised, merged and taxonomy assigned via the filterAndTrim (trimRight = c(0,10), maxN = 0, maxEE = 1, rm.phix=TRUE, truncQ = 20, trimLeft = c(19,20)), dada (pool = TRUE, USE_KMERS = FALSE, GAPLESS = FALSE, MIN_ABUNDANCE = 20,MIN_HAMMING = 5, KDIST_CUTOFF = 1, BAND_SIZE = -1), mergePairsand `assignTaxonomy` functions in the `R` package `dada2` (Callahan et al. 2016).

Multiple sequence alignment (MSA) was performed using the L-INS-i (Berger and Munson 1991) method in the software MAFFT (Katoh and Standley 2013). Subsequently a maximum likelihood phylogenetic tree was constructed in IQ-Tree (Nguyen et al. 2015) with 1000 bootstrap replicates and nucleotide substitution model selected by the built-inModelFinder, with Bayesian information criterion (BIC) minimisation as the model selection criteria.

### Monoculture growth

We tested the ability of each isolate to population to grow within 24 hours (OD600) in 96-well plates under 6 conditions: 100 µl of M9 minimal media with one of the 3 carbon sources or all 3 combined, with or without community supernatant (sterilised spent media from inoculating all 46 isolates together in the relevant growth media). A maximum optical density of 0.045 was chosen as the threshold for detectable growth, based on visual inspection of all growth curves.

### Statistical analyses

All statistical analyses were performed via Markvov chain Monte Carlo (MCMC) in the Julia programming environment (Bezanson et al. 2017) using the probabilistic programming language `Turing.jl` (Ge, Xu, and Ghahramani 2018): 4 chains, each consisting of 1000 iterations of the No U-Turns (NUTS) (Homan and Gelman 2014) sampler, with 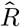 values < 1.01 for all parameters of interest. Credible intervals (CI) were calculated as the 2.5% and 97.5% quantiles for all parameter estimates and probability direction (PD) calculated for each contrast of interest as the proportion of MCMC samples in which the sign of the contrast (+/-) matches the sign of the point estimate of the contrast. Raw data and model estimates were visualised using Makie.jl (Danisch and Krumbiegel 2021).

### α-diversity

3 *α*-diversity metrics were analysed: strain richness, Pieou’s evenness index and Faith’s phylogenetic diversity (PD). In each case, two different modelling approaches were compared with model selection via Pareto-smoothed importance sampling leave one out crossvalidation (PSIS-LOO CV) (Vehtari, Gelman, and Gabry 2017): a *pooled* model where all replicate microcosms with a single carbon source are contrasted against all of those with multiple; a *hierarchical* model where each carbon source has its own regression coefficient, and the *α*-diversity of each multiple carbon source community is modelled as an additive combination of the model intercept and the regression coefficients of each carbon source it contains. For the hierarchical model, the parameter of interest is the mean of the distribution from which the regression coefficients are drawn. In each model, intercepts were given priors of normal(0,4), regression coefficients had priors of normal(0,3) (for hierarchical models this meant a normal(0,3) hyperprior over the mean of the distribution of regression coefficients). All variance parameters (including hyperparameters) had priors of exponential(1). Prior to analysis via linear models, Pielou’s evenness and Faiths PD were transformed onto the real number line by logit and log transformation respectively. Strain richness counts were analysed via Poisson models with a log link. For strain richness and Pielou’s evenness, the hierarchical model was selected, whereas for Faith’s PD the pooled model was the best model.

### β-diversity

4 *β*-diversity metrics were analysed: Bray-Curtis dissimilarity between strain proportions, Jaccard’s dissimilarity and both weighted and unweighted Uni-Frac. We used a mixed-model to estimate the expected dissimilarity between pairs of communities that do and do not have identical abiotic (carbon source) environments, dealing with non-independence of pairwise observation matrix entries by including a random effect for each community (using a Bayesian approach adapted by (Gompert et al. 2014) from an original maximum-likelihood based model developed by (Clarke, Rothery, and Raybould 2002)). We performed inference on each nutrient diversity treatment (single vs multiple carbon sources) separately. To analyse the impact of a shared carbon source we included 2 matrices of indicator variables in each model, *X*1_*i,j*_= 1 if the *i*^*th*^ and *j*^*th*^ samples were cultured on the same carbon source else = 0, *X*2_*i,j*_= 0 if the *i*^*th*^ and *j*^*th*^ samples were cultured on the same carbon source else = 1. Logistic transformed Bray-Curtis, and weighted and unweighted Uni-Frac values were analysed with linear models, while Jaccard’s dissimilarity was analysed with a binomial model with logit link, using the counts of shared strains and total strains for each pair of communities as *successes* and *trials*, described in detail in (Newbury, Kuijper, and Buckling 2023). All intercepts and random effects had priors of normal(0,4) and variance parameters had priors of exponential(1).

### Co-occurrence

The effect of phylogenetic distance (according to our maximum likelihood tree) on the cooccurrence relationships between strains was estimated using a similar mixed model approach to that used for *β*-diversity, but with co-occurrence affinity as the response variable (see (Newbury 2024)). Random effects and regression coefficients had priors of normal(0,10) which were wider priors than used in the other models in this work due to the lack of prior knowledge about how strong the relationship could plausibly be. The affinity of each pair of strains was also estimated, using the same MCMC methods as all other statistical models and plotted with the posterior median as the point estimate and the interquartile range of the posterior distribution shown as error bars.

## Notes

### Competing Interest Statement

The authors have declared no competing interest.

