## Supplemental Figure 1 for "Diversity *Between* but Not *Within* Microbial Communities Increases With the Diversity of Supplied Nutrients"

### Beta diversity without *Flavobacterium sp.*

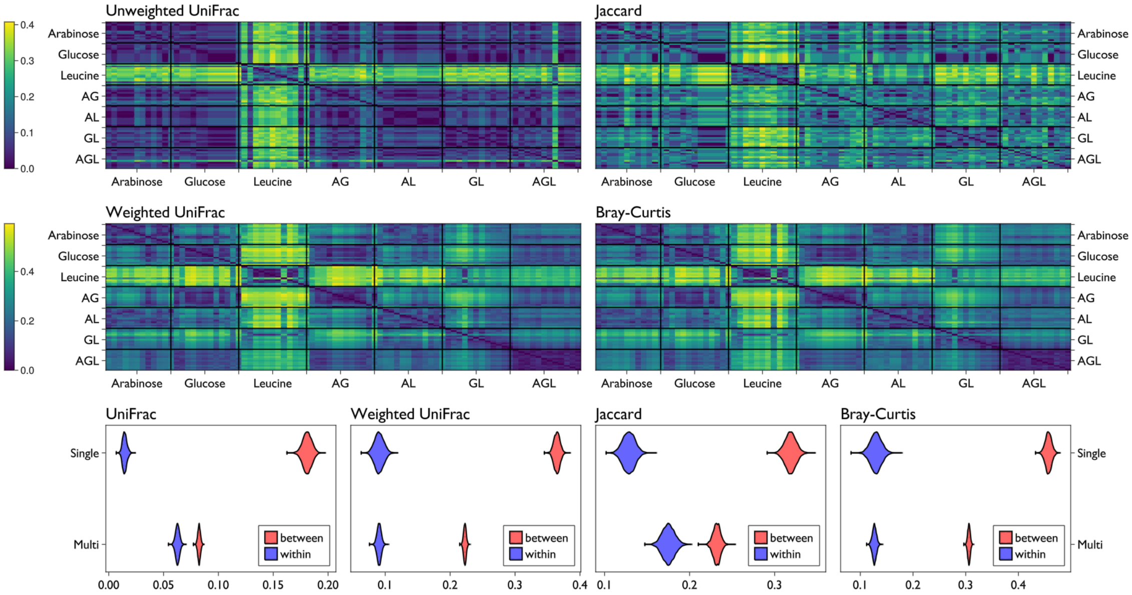


**Supplemental Figure 1**. Beta diversity estimates as in the main text but with the removal of the rare but phylogenetically distinct *Flavobacterium sp.*
